# Precision culturomics enabled by unlabeled single-cell morphology and Raman spectra

**DOI:** 10.1101/2025.06.27.661433

**Authors:** Qiaoxing Liang, Xihong Lan, Jiayi Wu, Wei Wei, Lili Li, Xiaofang Tang, Guoping Zhao, Ruidong Guo, Huijue Jia

## Abstract

Selective enrichment of target bacteria from complex communities such as the human microbiome has remained a challenge. Here we report a solution based on morphology, Raman spectrometry and the Laser-Induced Forward Transfer technology, which works at microbial single cells, many generations before they appear as colonies. We develop a machine learning-based framework that enables species-level targeted sorting of single microbial cells from complex microbiome. We illustrate the utility of this approach in selecting for or against specific bacteria in fecal microbiome samples and the potential for quantifying the molecules expressed based on Raman spectra. Analysis of single-cell cultured genomes reveals that brief antibiotic use drives both pre-existing resistance and *de novo* mutations in the transpeptidase or efflux pumps of gut commensals, along with convergent evolution between different species. Our precision culturomics method should enable detailed morphological, metabolic, and genomic insights into variations in microbial phenotypes at the single-cell level for microbiome studies.

## Introduction

Functional characterization of the microbiome, and better standardization of commonly used animal models require defined isolates of commensal microbes. Despite recent progress in culturomics, bacterial isolates obtained by plating and colony-picking tend to retrieve bacteria that are abundant in the sample^1,2^. Image analyses of the colonies could allow some selection, but the enrichment often did not exceed 50% at the genus level^3^. Although no longer deemed unculturable, incubation time for commensal microbes can take weeks or months^4^, and the plates become more suboptimal with time, due to dehydration, and conversion of nutrients. To be able to selectively retrieve specific taxa at the single-cell stage would therefore be time-saving, resource-efficient, and ensures recovery of relevant microbes.

Laser-Induced Forward Transfer (LIFT) is a straight-forward way of manipulating microparticles^5^. With a laser beam pushing on the matrix, LIFT is ideally suited for picking particles in the size range of bacteria, and the ejected cells can be collected in a short distance. LIFT is rarely combined with Raman spectrometry, which is increasingly studied for rapid identification of bacteria such as methicillin-resistant *Staphylococcus aureus* (MRSA)^6^. Labeled approaches, such as isotope probing or substrate analogue probing, fail to provide sufficient information for taxonomic discrimination and may negatively impact cell activity. In contrast, label-free properties, including cell morphology detected through optical microscopy, and chemical composition detected through Raman spectrometry, could offer valuable insights for taxonomic identification at the single-cell level. Moreover, LIFT could retain spatial information of the retrieved microbes in the microbiome sample.

Here we achieve single-cell culturomics from LIFT, and pick the single cells according to morphology and Raman spectra (**Fig. 1a**). The approach consists of three key procedures outlined as follows: (1) collection of single-cell morphological and Raman spectral data from microbiome samples, (2) identification of microbial cells belonging to target taxa using a machine learning-based framework, and (3) cell sorting via LIFT and culturing from single cells. The approach enables identification of specific microbial species from complex microbiome samples. We demonstrated the utility of this approach by selectively culturing for or against specific species from the human gut microbiota. This in situ single-cell omics^7^ approach is poised to greatly facilitate pharmaceutical developments in the host-associated microbiome and further our understanding of natural communities. Future development of more specific applications may also include routine quality examination of Live Biotherapeutic products, safety monitoring of fecal transplants, etc.

**Figure 1.**
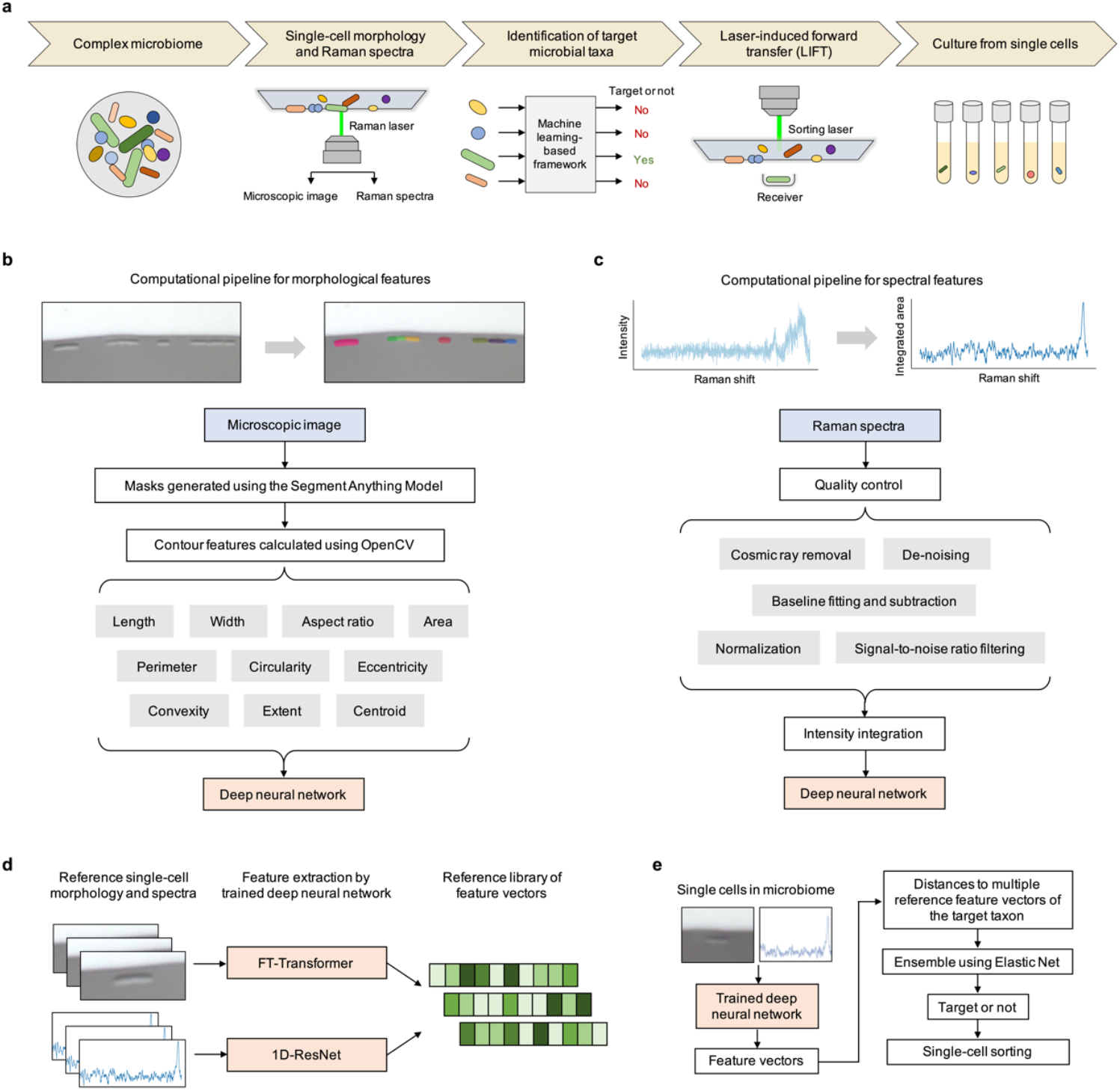
Overview of the precision single-cell culturomics. (**a**) Schematic for the method. Single-cell morphological and Raman spectra data of complex microbiome samples were collected by the inverted microscope. A machine learning-based framework was used to identify whether these cells belonged to target microbial taxa. The microbial cells of the desired taxa were sorted by LIFT on the commercially available PRECI-SCS equipment, and cultured in tubes in an incubator set at the desired conditions. (**b**) Computational pipeline for morphological features. (**c**) Computational pipeline for Raman spectral features. (**d**) Generation of a single-cell reference library of feature vectors with known taxonomic labels using a trained deep neural network. Features were extracted from single cell morphological and spectral data using FT-Transformer and 1D-ResNet respectively. (**e**) Feature vectors of single cells in microbiome samples were extracted using a trained deep neural network. The distances of these vectors to multiple reference feature vectors for a specific taxon were input into an elastic net to determine whether the cells belonged to that taxon. Cells identified as belonging to the target taxa were then sorted for culturing.

## Results

### Single-cell live culture from microliter samples

With precision culturomics in mind, we set out to test whether bacteria are still alive after the single-cell ejection by LIFT using a commercially available device (**Supplementary Fig. 1**). One advantage for this method would be the microliter amounts of precious samples required. We performed Caesarean section for mice, spotted the amniotic fluid onto the aluminum-coated quartz slide, and LIFT-picked the bacterial cells before complete drying of the sample (**Supplementary Fig. 1**). After prolonged anaerobic incubation for two weeks, high-density populations emerged in some of the wells. Among the isolates were interesting species such as *Oceanobacillus picturae*, which could produce ectoine for skin health^8^, and *Pseudomonas* sp. known to be present in the upper reproductive tract of humans and could metabolize estrogen^9–12^. Other bacterial species included *Klebsiella pneumoniae* and *Enterococcus lactis*. We searched for their closely related genomes in the human microbiome using the MGnify genome collections of the human gut, oral, and vaginal microbiomes^13^. The most closely related genomes within the same species (Mash distance < 0.05) were identified in the gut for *O. picturae, K. pneumoniae* and *E. lactis*.

### Harnessing microbial single-cell morphology for species-level discrimination

In order to identify specific taxa before culture, we collected single-cell morphological and Raman spectral data from isolated strains of 25 bacterial species across 14 genera commonly found in the human microbiome to train machine-learning models. Using this data resource, which links single-cell morphology and spectra to taxonomy, we first examined whether single-cell morphology can help distinguish bacteria at the species level. The morphological data of single cells were measured from microscopic images (**Fig. 1b**). We developed a cell contour analysis pipeline that performs image segmentation to generate masks for cells in an image, utilizing the Segment Anything Model. These masks are then processed using the computer vision tool OpenCV to calculate ten morphological features related to cell size and shape. We performed principal component analysis using the ten morphological features measured from microscopic images. As expected, spherical bacteria, such as *Veillonella, Staphylococcus, Lactococcus*, and *Streptococcus*, tended to be closer to each other, while rod-shaped bacteria like *Bacillus, Bifidobacterium*, and *Clostridium* showed a similar tendency (**Fig. 2a**). Cell length and width were the most dominant features for principal components 1 and 2, respectively (**Fig. 2b**). In addition to length, perimeter, circularity, and aspect ratio made significant contributions to principal component 1, which was the primary axis along which cells from different taxa were separated and accounted for 53.38% of the morphological variance.

**Figure 2.**
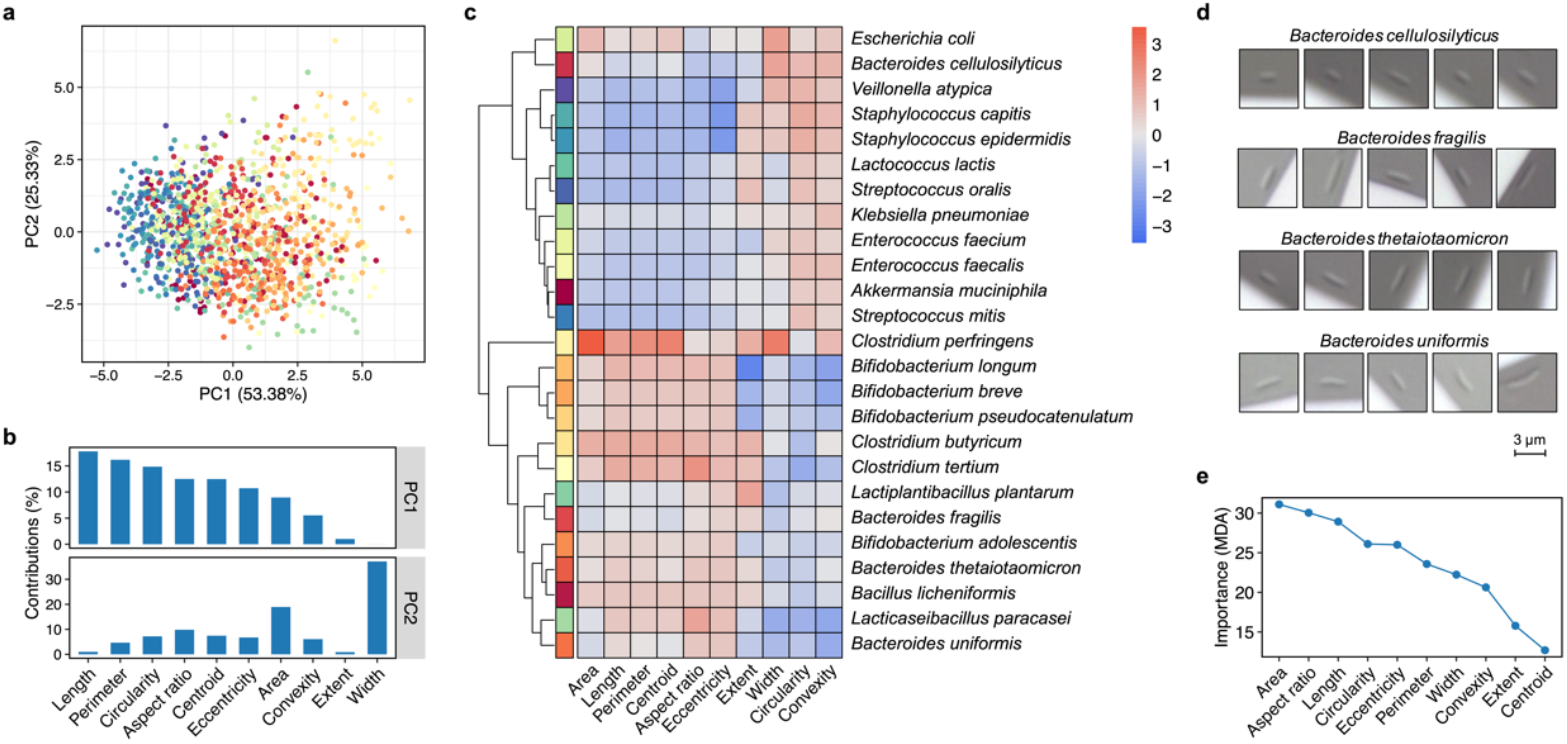
Morphological features of microbial single cells contribute to species-level discrimination. (**a**) PCA ordination of microbial single cells based on morphological features. Colors indicate species-level taxonomy. (**b**) Informative features in PCA analysis of single-cell morphology. Contributions of morphological features to principal component 1 and principal component 2 were showed. (**c**) Heatmap of median *z* scores of morphological features across different microbial species. Colors for each row on the left correspond to colors of the dots in the PCA plot. (**d**) Examples of single-cell images of different *Bacteroides* species. (**e**) Importance scores of morphological features measured as mean decrease in accuracy (MDA) in a random forest model for multi-species classification.

Single cells from different species exhibited diverse morphological patterns (**Fig. 2c; Supplementary Fig. 2**). Moreover, cells from different species within a genus can vary significantly in their morphology. For example, *Bacteroides cellulosilyticus* was relatively short and almost oval in shape, while *Bacteroides uniformis* and *Bacteroides thetaiotaomicron* were much longer and closer to typical rod-shaped bacteria (**Fig. 2d;** *P* < 2.2×10^−16^). While *Clostridium butyricum* and *Clostridium tertium* were clustered together, *C. butyricum* was significantly wider than *C. tertium* (*P* = 2.7×10^−11^), and *C. perfringens* was separated from them in the clustering diagram. In order to gain insight into the relative contribution of different morphological features to species-level classification, a random forest multi-class classification model was trained using the ten morphological features. The importance scores, measured as the mean decrease in accuracy, were greater than 10% for all the features. The most important feature was area, followed by aspect ratio, length, circularity, eccentricity, perimeter, and width. In conclusion, the morphology of a bacterial single cell contains a considerable amount of information that is useful for discrimination at the species level.

### Linking single-cell phenotypic variation reflected in morphology and spectra

The analysis of single-cell morphological features above revealed that cells of the same species displayed significant morphological heterogeneity. In addition to morphology, intra-species variation at the cell level can also be reflected in chemical composition, which can be detected through Raman spectroscopy. To achieve relative quantification of molecules from Raman spectra, we collected spectra of pure compounds and created a library of important cellular components, including DNA and RNA bases, amino acids, primary metabolites, cell wall components, and other key molecules (**Supplementary Fig. 3**). We developed a method to fit a cellular spectrum using the spectra of multiple pure compounds, thereby enabling an estimate of quantification to be obtained based on the weights assigned to each compound. Quality-controlled Raman spectra are converted from intensity to integrated area (± 10 cm^-1^) along the Raman shift (**Fig. 1c**).

To assess the feasibility of this method of molecular characterization from Raman spectra, we collected Raman spectra from *Bacillus licheniformis* cells in three states: (1) vegetative cells cultured in Columbia Broth as a control, (2) pre-sporulation vegetative cells cultured in 10% Columbia Broth with pH adjusted to 4.0 by citric acid, and (3) spores formed following the citric acid treatment. Vegetative cells and spores were selected visually. We found that the spectra of spores were significantly different from those of vegetative cells (**Fig. 3a,b**). The molecules that exhibited differences among the three states were identified based on the results of spectrum fitting (**Fig. 3c**). Dipicolinic acid, a crucial component in bacterial spores that enhances resistance to environmental stresses^14^, was most abundant in spores, followed by pre-sporulation cells, and least abundant in control cells (*P* < 1.0×10^−8^). In spores, the outermost layer is no longer the peptidoglycan cell wall, but the spore coat, which is primarily composed of proteins^15^. Accordantly, we detected a significantly weaker signal for peptidoglycan in spores than in vegetative cells, along with a stronger signal for phenylalanine, an amino acid commonly used as a marker for protein analysis in Raman spectroscopy (*P* < 2.2×10^−16^). Arginine, which plays an important role in bacterial endospore formation through trafficking between the forespore and the mother cell^16^, was significantly more abundant in pre-sporulation cells than in control cells (*P* = 1.7×10^−6^). Vegetative cells actively utilize citric acid in metabolic processes such as the Krebs cycle, which are largely inactive in dormant spores. Consistently, citric acid was more abundant in vegetative cells than in spores (*P* < 2.2×10^−16^). These results indicate that such a data-analysis approach can be used for quantifying molecules of interest from single-cell Raman spectra.

**Figure 3.**
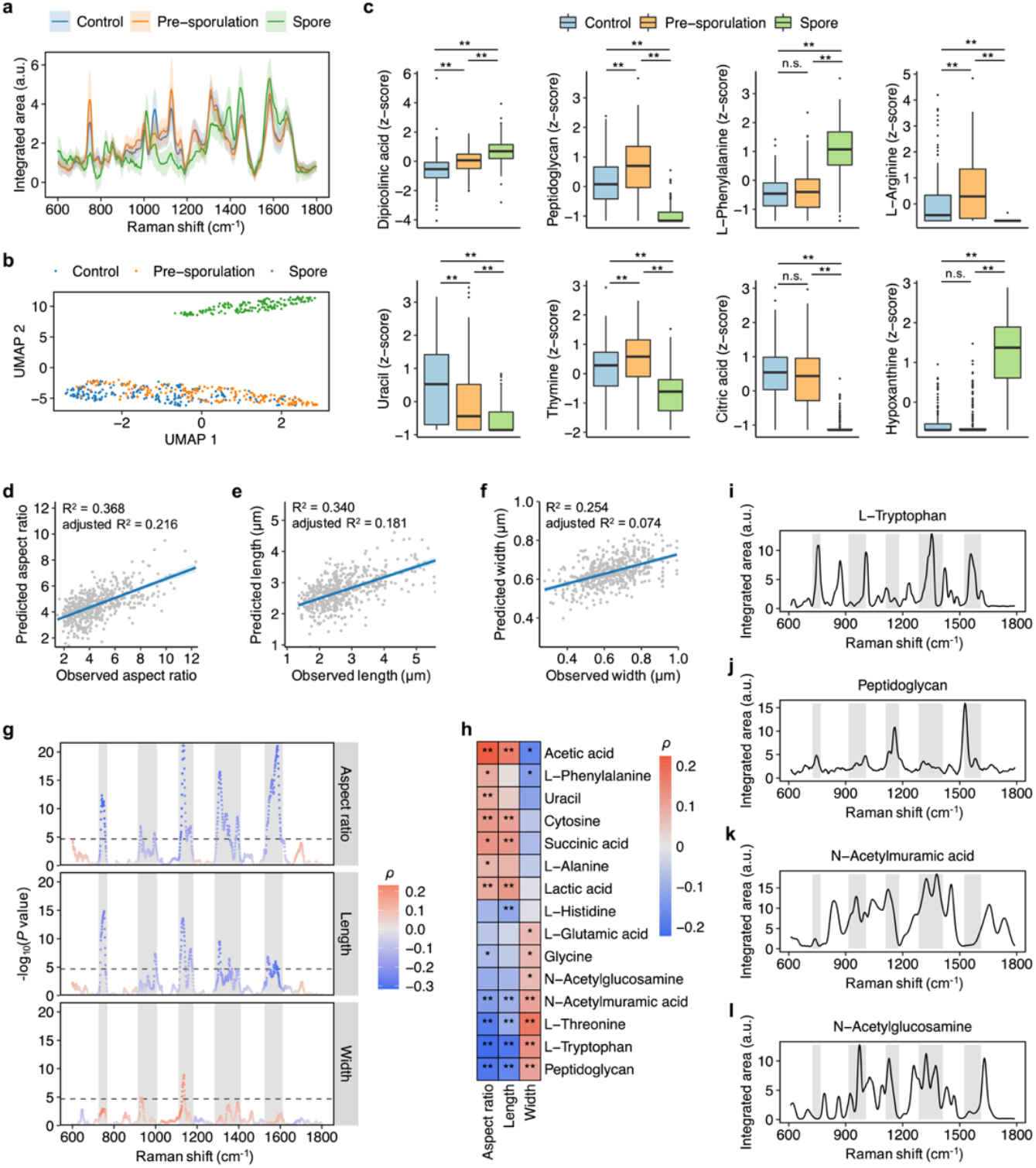
Associations between single-cell morphology and Raman spectra. (**a**) Raman spectra of *B. licheniformis* vegetative cells in normal or sporulation broth and spores. Data were represented as ± 10 cm^-1^ integrated area for each Raman shift (mean ± standard deviation). (**b**) UMAP embeddings of Raman spectra colored by the three states of *B. licheniformis* cells. (**c**) Examples of molecules that show differences in quantification among the three states of *B. licheniformis* cells (FDR \**P* < 0.05, \*\**P* < 0.01). Data were shown as *z* scores across cells. (**d**-**f**) Measured (*x* axis) and predicted (*y* axis) single-cell aspect ratio (d), length (e), and width (f) based on Raman spectra. (**g**) Correlations of Raman spectra with single-cell aspect ratio, length, and width. The color of the dots represents the Spearman correlation coefficient (ρ). The dashed horizontal line indicates a significance threshold of *P* < 2.08×10^−5^ (0.05 / the number of Raman shifts). The gray shading indicates intervals that show significant associations. (**h**) Correlations of molecular quantifications estimated by Raman spectra with single-cell aspect ratio, length, and width (\**P* < 0.05, \*\**P* < 0.01). (**i**-**l**) Raman spectra of molecules that may contribute to the correlations between single-cell morphology and Raman spectra. The gray shading corresponds to the intervals in (g).

Simultaneous collection of morphological and Raman spectral data provides an opportunity to characterize single-cell phenotypic variation, linking morphological and molecular heterogeneity. Using data from cells of the same *Bacillus licheniformis* strain, we investigated associations between single-cell morphology and spectra. To assess the extent to which morphological variance can be explained by Raman spectra, we fitted the morphological features using multivariable linear regression models with the top 100 principal components of spectra. The coefficient of determination was highest for aspect ratio (R^2^ = 0.368), followed by length (R^2^ = 0.340) and width (R^2^ = 0.254). The predicted and observed values of morphological features were significantly correlated (*P* < 2.2×10^−16^; **Fig. 3d-f**). We performed a correlation analysis between these features and the integrated area for each Raman shift, identifying five spectral regions that were negatively correlated with aspect ratio and length (Spearman correlation, Bonferroni-corrected *P* < 0.05). These regions included 736-758 cm^-1^, 925-997 cm^-1^, 1121-1172 cm^-1^, 1294-1400 cm^-1^, and 1534-1602 cm^-1^, among which the range 1121-1172 cm^-1^ was also positively correlated with width (**Fig. 3g**). We further fitted the cellular spectra to those of pure compounds and identified correlations between specific molecules and morphological features (**Fig. 3h**). For example, tryptophan was negatively correlated with aspect ratio (FDR = 9.1×10^−7^) and length (FDR = 1.4×10^−8^), and may partially account for the negative correlations in the spectral regions of 736-758 cm^-1^, 1294-1400 cm^-1^, and 1534-1602 cm^-1^ (**Fig. 3i**). Peptidoglycan, the primary component of the cell wall, also showed negative correlations with aspect ratio (FDR = 2.0×10^−8^) and length (FDR = 1.9×10^−6^). Peptidoglycan and its building blocks, N-Acetylmuramic acid and N-Acetylglucosamine, may partially account for the negative correlations in 1121-1172 cm^-1^ and 925-997 cm^-1^ (**Fig. 3j-l**). Thus, within the same strain, bacterial single-cell morphological features such as cell length and aspect ratio are partly reflected in the cells’ Raman spectra.

### Species-level classification models based on morphological and spectral features

The machine learning-based framework consists of two parts: (1) extracting feature vectors that minimize distances between cells of the same taxon while maximizing distances between cells of different taxa, using single-cell morphology and Raman spectra with a deep neural network, and (2) determining whether a cell belongs to the desired taxon by comparing the distance between its feature vector and those of reference cells. Feature vectors of reference cells are calculated using the trained network and stored in a reference library (**Fig. 1d**). Distances to multiple reference cells of a specific taxon are integrated using multivariable logistic regression to predict whether a cell belongs to that taxon (**Fig. 1e**).

We adopted contrastive learning based on a Siamese network to extract feature vectors from microbial single-cell morphological and Raman spectral data (**Fig. 4a**). The microbial cells used for training are organized into positive pairs (cells from the same taxon) and negative pairs (cells from different taxa). The two cells are mapped into an embedding space using an identical deep neural network. Contrastive loss encourages the model to learn an embedding space where cells from the same taxon are close together, while cells from different taxa are far apart. After training, the cells used for training were fed into the trained network to extract feature vectors, which constitute a reference library with each taxon represented by multiple cells (**Fig. 4b**).

**Figure 4.**
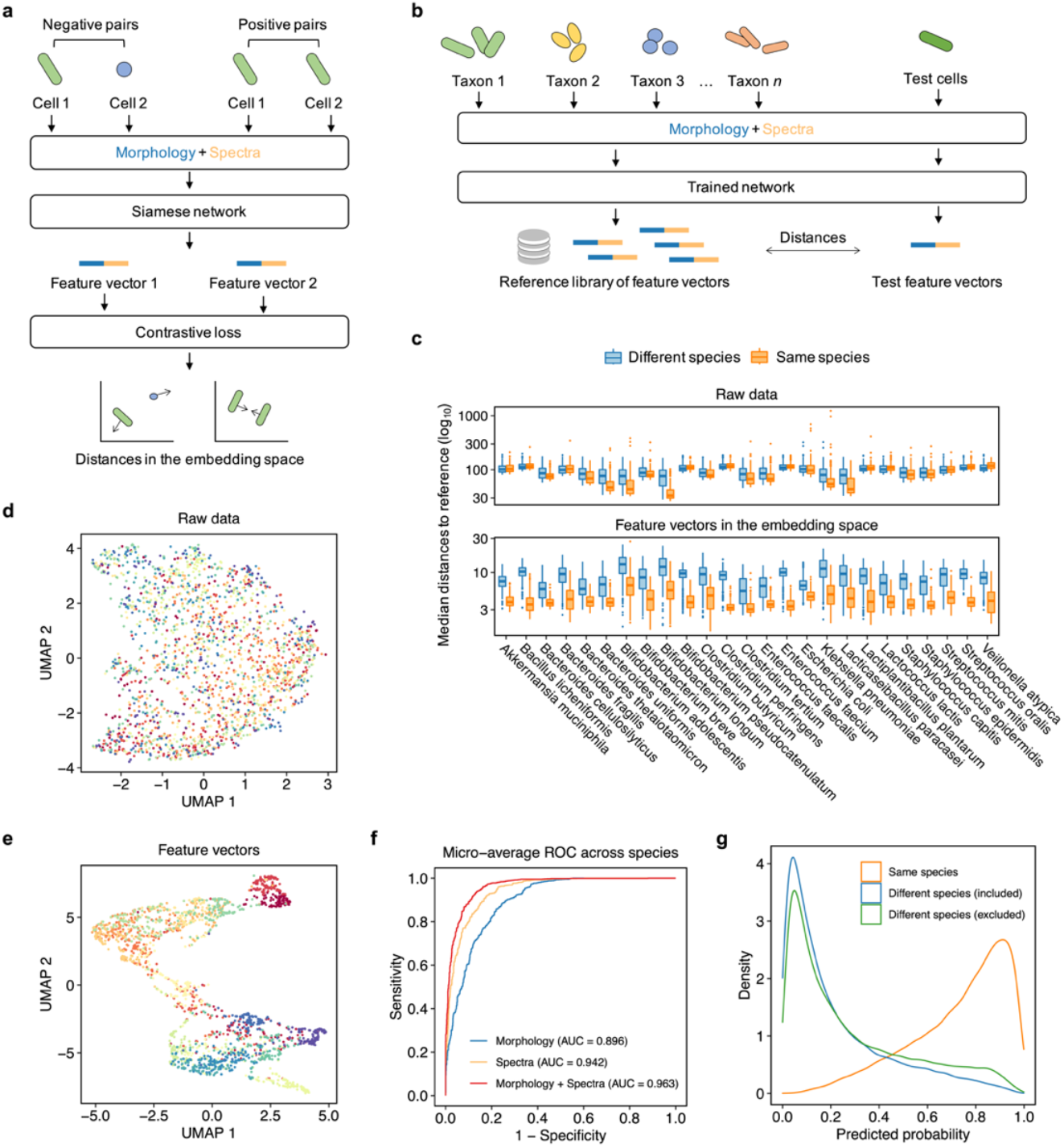
The machine learning-based framework for taxonomic identification. (**a**) Schematic for the contrastive representation learning of single-cell morphological and Raman spectral data. Single cells were organized into positive and negative pairs, where positive pairs consist of cells from the same taxon, and negative pairs consist of cells from different taxa. The morphology and spectra of two cells were processed by Siamese network. Contrastive loss was applied to minimize distances for the same taxa and maximize distances for different taxa in an embedding space. (**b**) Schematic for cell matching based on features extracted by a deep neural network. A reference library of feature vectors with taxonomic labels was created. Distances between the feature vector of a test cell and those of multiple reference cells were then calculated. (**c**) Median Euclidean distances of raw data (concatenation of morphological and spectral data) or feature vectors in an embedding space between test cells of different or the same species and reference cells. (**d, e**) UMAP embeddings of raw data (d) and feature vectors (e). Colors indicate species-level taxonomy. (**f**) Receiver Operating Characteristic (ROC) curves for elastic net predicting target species based solely on morphology, Raman spectra, or a combination of both morphology and spectra. (**g**) Distribution of the predicted probabilities of being a target species for test cells from the same or different species that were included in or excluded from the training sets.

Similarly, feature vectors are extracted from the morphological and Raman spectral data of new cells encountered in microbiome samples. The distances of the feature vectors between multiple reference cells of a taxon and a new cell are used as predictors in a regularized multiple linear regression model, such as elastic net, to predict whether the new cell belongs to that taxon.

We constructed a Siamese network using data from 25 bacterial species commonly found in the human microbiome. Feature vectors for morphological and spectral data were extracted using FT-Transformer and ResNet, respectively, and then concatenated. The network was trained at the species level, encouraging cells from the same species to be close together while cells from different species were pushed farther apart. For each species, we calculated the Euclidean distances between reference cells and test cells from the same or different species (**Fig. 4c**). We observed that the distances calculated from feature vectors in the embedding space are significantly lower for reference cells and test cells of the same species compared to those of different species. In contrast, distances calculated from raw model inputs for the two groups are less distinguishable. Consistent with these results, the UMAP visualization based on raw data showed cells intermingled across species, whereas the UMAP visualization based on feature vectors revealed clusters for species (**Fig. 4d,e**).

To evaluate the utility of single-cell morphology and Raman spectra, as well as the advantages of integrating both data in taxonomy prediction, we also constructed deep neural network that used either morphological or spectral data as input. We trained a separate elastic net model for each species to perform binary classification based on distances derived from the three types of inputs. Even for models based exclusively on morphological data, the area under the receiver operating characteristic curve (AUC-ROC) approached 0.9 for the majority of species (**Fig. 4f**; **Supplementary Fig. 4**). The AUC-ROC of models utilizing both data types were significantly higher than those of models based solely on morphology (Wilcoxon signed-rank test, *P* = 2.3×10^−5^) or on spectra (*P* = 1.5×10^−2^). For some species, models based solely on morphology or spectra achieved performance comparable to that of models integrating both data types. These results demonstrated that single-cell morphological and Raman spectral data are both effective in discriminating between different species. In general, integrating both data types can enhance model performance.

We further constructed models at the genus level by labeling microbial cells from the same genus as positive pairs. The deep neural network successfully learned a feature embedding space in which cells formed distinct clusters corresponding to different genera (**Supplementary Fig. 5**). Consistent with the species-level elastic net models, the genus-level models based on distances derived from both morphology and spectra outperformed those based solely on morphology (*P* = 1.7×10^−3^) and spectra (*P* = 1.7×10^−3^) (**Supplementary Fig. 6**). The overall performance assessed by the micro-average AUC-ROC across taxa was slightly lower in genus-level models compared to species-level models, which may be due to the increased intra-class variation. Nonetheless, for models using both morphological and spectral data, both the species-level and genus-level models achieved a micro-average AUC-ROC greater than 0.9.

Given the high diversity of the microbiome, models should avoid misclassifying cells from taxa that were not included during training as target cells. To evaluate this, we constructed leave-one-out models that excluded one species or genus from training at a time and made predictions for cells from the excluded taxa. We observed that the distribution of predicted probabilities for these cells was similar to that of cells from non-target taxa included in the training (**Fig. 4g**; **Supplementary Fig. 5**). This attribute allows for the establishment of a probability threshold to effectively separate target and non-target cells based on benchmark results from restricted datasets, rather than the entire microbiome.

### Culturomics from complex microbiome samples with selection

To demonstrate the utility of our experimental and computational approach in complex microbiomes, we performed single-cell cultures from fecal samples. The LIFT module successfully isolated individual bacterial cells without disturbing neighboring cells through laser energy-tuned ejection size adjustment (0.5-10 μm, **Supplementary Figs. 1**,**7**). For fresh fecal samples, single-cell viability rate determined by broth turbidity (>10^8^ cfu/ml according to *Escherichia coli*) within three weeks averaged 62.50%. This approach captured diverse bacteria, including abundant species and those undetectable by marker gene-based tools like MetaPhlAn 4^17^ (**Fig. 5a**). To excluding the effect of cultivability limitations on viability, we conducted an overnight pre-culture of fecal samples under conditions identical to single-cell cultures. The pre-culturing step increased the viability rate to 91.67%. We found that 71.91% of the resulting cultures were *E. coli* (**Fig. 5b**). This situation represents a typical challenge in traditional culturomics: fast-growing and abundant bacteria in the samples are more likely to be cultured and thus dominate the colonies^1,2^. To demonstrate machine learning applications in culturomics depleted for specific bacteria, we implemented *E. coli* negative selection in pre-cultured fecal samples and achieved 92.86% non-*E. coli* recovery. At a prediction probability threshold > 0.3, 92.31% of the predicted *E. coli* cells were true positives. For the cells misclassified as *E. coli*, target models showed higher prediction probabilities than the *E. coli* model, suggesting the feasibility of recalling target species in negative selection. Average nucleotide identity (ANI) analysis of identified *E. coli* cells revealed different strains compared to the reference strain for training, confirming model generalizability to untrained strains (**Fig. 5c**).

**Figure 5.**
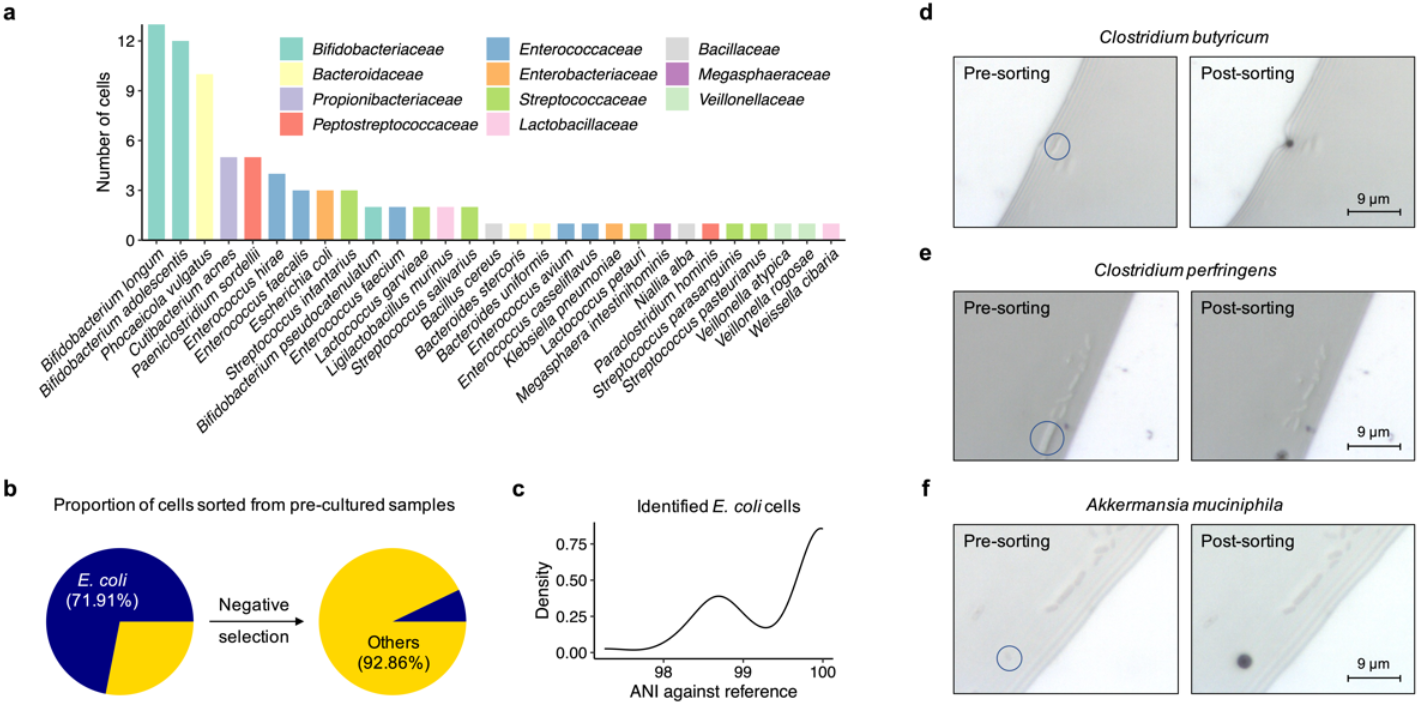
Precision culturomics for the gut microbiota. (**a**) Number of isolates obtained by single-cell sorting from gut microbiota samples for different bacterial species. (**b**) Proportion of bacterial cells sorted from pre-culture fecal samples, with and without negative selection against *E. coli*. (**c**) Distribution of ANI of identified *E. coli* cells against the reference strain. (**d**-**f**) Microscopic fields before and after sorting for *C. butyricum* (d), *C. perfringens* (e), and *A. muciniphila* (f).

Having demonstrated the utility of our approach in selecting against unwanted dominant taxa, we next applied our method to the enriched cultivation of specific bacterial species. As a proof of concept, we selectively isolated beneficial *Clostridium* species while excluding pathogenic ones. *C. butyricum*, an approved Live Biotherapeutic Product before the revived interested in microbiome, is renowned for its ability to produce butyrate, is beneficial for health as it supports the integrity of the gut barrier, regulates inflammation, aids in restoring gut microbiota after antibiotic use, and may have therapeutic potential in managing conditions like diabetes and gastrointestinal infections^18^. In contrast, pathogenic *C. perfringens* produces toxins causing tissue damage and immune dysregulation^19^. Using selection criteria requiring *C. butyricum* model probability > 0.6 and higher than other *Clostridium* species, we successfully isolated two *C. butyricum* cultures. Consistent with the marked morphological differences between *C. butyricum* and *C. perfringens*, our models correctly distinguished them using only morphological data (**Fig. 5d,e**). We further isolated a culture of *C. tertium*, which was morphologically similar to *C. butyricum*, with morphology models assigning probabilities of 0.770 (*C. tertium*), 0.010 (*C. butyricum*), and 0.002 (*C. perfringens*), confirming morphological discriminative power. To showcase the capability of our method for rare taxa isolation, we selectively cultured for *Akkermansia muciniphila*, a beneficial gut species whose abundance was 0.0017% in the sample used, and successfully yielded a culture (**Fig. 5f**). In conclusion, our method can be used to culture either for or against specific bacteria from complex microbiome samples.

### Resistance of human gut commensal species driven by antibiotic exposure

The impact of short-term antibiotic use on beneficial gut bacteria has become a critical concern in clinical medicine^20^. Sequencing-based approaches for studying antibiotic effects face computational challenges, such as resolving genetic variants into strains, analyzing species with low sequencing depth, and distinguishing low-frequency mutations from sequencing errors. To illustrate how culturomics can overcome these limitations, we isolated single cells of two beneficial *Bifidobacterium* species—*B. longum* and *B. adolescentis*—from a human volunteer (H1) who received a 5-day course of oral cefuroxime for non-intestinal surgery. *B. longum* has been reported to reduce throat pain and symptoms like runny nose and cough in children, alleviate allergic rhinitis and intermittent asthma, improving quality of life^21^. *B. adolescentis* is a species that appeared earlier in formula-fed infants compared to breastfed infants, and recently implicated with antibody counts after COVID-19 vaccination^22,23^, and increased healthy lifespan in multiple model organisms^24^. Fecal samples were collected before cefuroxime administration (with no antibiotic use in the preceding year) and five weeks after treatment ended.

Using a threshold of ANI > 99.5%, we identified two *B. longum* strains in H1, with a mean ANI of 98.65% between them (**Fig. 6a,b**). We first analyzed penicillin-binding protein 2 (PBP2), the direct target of cefuroxime in *B. longum*, and found that each strain carried a distinct PBP2 variant differing at three amino acid positions. For clarity, we designated the two variants as v1 (430H, 546V, 590G) and v2 (430R, 546A, 590S). Based on cell counts, the strain carrying v1 was dominant over the one carrying v2. As a control, we isolated *B. longum* from a volunteer (H2) who reported no antibiotic use in the preceding three years. *B. longum* from H2 represented another strain with a mean ANI of 98.54% relative to H1 strains and harbored PBP2 variant v3 (168A, 430R, 546A, 568S, 590S). To assess functional implications, we predicted the structures of all three variants and performed cefuroxime docking (**Fig. 6c**). The mutations distinguishing v1 and v2 localized to PBP2’s transpeptidase domain, where cefuroxime binds (**Fig. 6d**). Affinity prediction and docking scoring suggested that v2 had reduced binding capacity compared to v1 and v3 (**Fig. 6e,f**). Notably, both v1 and v2 strains persisted before and after cefuroxime exposure, but the v2 population exhibited a trend toward increase after exposure (Fisher’s exact test, *P* = 0.1), suggesting evolution of pre-existing resistance at the population level.

**Figure 6.**
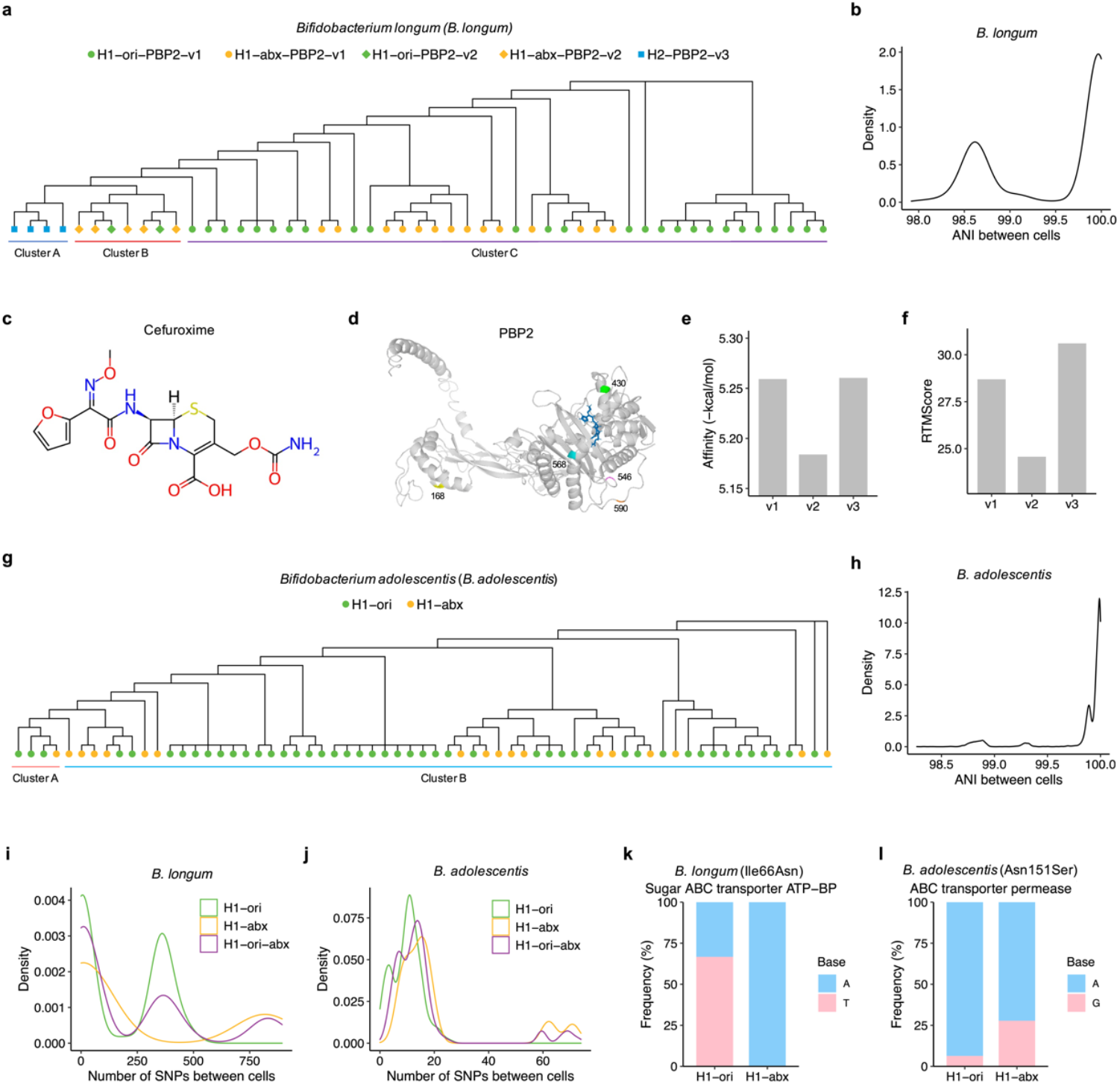
The impact of brief antibiotic use on specific gut bacteria. (**a**) Phylogenetic tree constructed for different cells of *B. longum*. Cells are grouped into clusters using a threshold of ANI > 99.5%. The shape of each node indicates variants of the PBP2 protein sequences, while the color indicates whether the cell was obtained before (ori) or after (abx) antibiotic exposure. (**b**) Distribution of ANI between different cells of *B. longum*. (**c**) Chemical structure of cefuroxime obtained from the ChEMBL database. (**d**) Structure of PBP2 protein bound to cefuroxime. (**e**) Predicted binding affinities between three variants of PBP2 protein and cefuroxime. (**f**) Scores generated using RTMScore for the optimal protein-ligand binding pose of three variants of PBP2. (**g**) Phylogenetic tree for different cells of *B. adolescentis*. (**h**) Distribution of ANI between different cells of *B. adolescentis*. (**i, j**) Number of SNPs among different cells obtained before (ori) or after (abx) antibiotic exposure, as well as between cells obtained before and after exposure (ori-abx) for *B. longum* (i) and *B. adolescentis* (j). (**k**) Allele frequency of a missense variant in the gene encoding the sugar ABC transporter ATP-binding protein (ATP-BP) of *B. longum*. (**l**) Allele frequency of a missense variant in the gene encoding the ABC transporter permease of *B. adolescentis*.

Similar to *B. longum*, individual H1 harbored two *B. adolescentis* strains, one of which was dominant, with both strains persisting before and after cefuroxime exposure (**Fig. 6g**). The dominant strain showed evidence of sub-strain diversification (**Fig. 6h**). In contrast to *B. longum*, all *B. adolescentis* cells shared identical penicillin-binding protein sequences, underscoring the role of pre-existing resistance mechanisms. We next investigated convergent evolution between the two species. While the number of pairwise SNPs between cells was generally lower in *B. adolescentis* than in *B. longum*, both species showed an increase in pairwise SNPs after exposure (Wilcoxon rank sum test, *P* < 0.05; **Fig. 6i,j**), suggesting *de novo* mutagenesis. To rule out strain proportion effects, we analyzed allele frequency changes within the same sub-strain before and after exposure. In *B. longum*, we identified a missense variant in the gene encoding sugar ABC transporter ATP-binding protein, with the T-allele depleted after exposure (Fisher’s exact test, FDR < 0.05; **Fig. 6k**). The ATP-binding protein and the permease are core components of the ABC transporter system, which is essential for sugar import but can also act as an antibiotic efflux pumps. Intriguingly, *B. adolescentis* exhibited a frequency shift in a missense variant within its ABC transporter permease gene (*P* = 0.03; **Fig. 6l**). Together, these findings reveal the distinct yet convergent adaptation strategies employed by beneficial gut bacterial species under short-term antibiotic stress.

## Discussion

To achieve precision culturomics at the single-cell level, we have developed a system that consists of collecting the single-cell morphology and Raman spectra, identifying targeted taxa using a machine learning-based framework, sorting single cells using LIFT, and subsequently culturing the sorted single cells. We demonstrated that this system enables the unlabeled sorting and culturing of desired taxa from complex microbiome samples.

The microbiome typically contains thousands of species, which could be resource-intensive to be all included in training datasets. This situation can lead to out-of-distribution (OOD) challenges, which are particularly problematic for multi-class classification strategies. Specifically, models may incorrectly classify OOD samples as one of the known classes, resulting in erroneous predictions. The machine learning strategy presented in this study for taxonomy prediction consists of two stages: feature vector extraction from single-cell morphological and spectral data using deep neural networks; the integration of distances between feature vectors and multiple reference cells from a specific taxon to determine whether a cell belongs to that taxon. This approach reframes the multi-class classification problem as reference database matching followed by binary classification. In addition to enhancing interpretability, this method also circumvents the challenges associated with confidence calibration for the model’s output probabilities. Furthermore, considering the scalability of contrastive learning—demonstrated in face recognition tasks involving datasets with tens of thousands of individuals and only a few samples per individual—it is feasible to achieve few-shot learning for single cells in the future. A promising starting point would be the construction of a large-scale database of single-cell morphological and spectral data.

We achieved species-level identification of bacterial cells using unlabeled features. Interestingly, the species-level models generally outperformed the genus-level models, possibly due to lower intra-class variation, and demonstrating the high resolution of our single-cell approach. This finding suggests future applications of our approach in fine-grained scenarios, such as strain-level identification. Additionally, we demonstrated the potential for molecular quantification using Raman spectra. These results imply the possibility of extending precision culturomics from specific microbial taxa to strains with particular metabolic functions, all without the need for labels.

Although we primarily focused on the human microbiome in this study, the system we presented can be applied to microbiome samples from various habitats. The preservation of spatial information can be further explored in combination with host data. The system utilizes commercially available equipment, and we have implemented the proposed machine learning-based framework as an easy-to-use Python package. We believe this system is ready to be adopted in laboratories around the world to facilitate microbiome research.

## Methods

### Microbiome samples

Amniotic fluid was collected from two pregnant mice using a sterile syringe inserted into the amniotic sac under direct visualization to avoid damaging surrounding tissues. Aseptic technique was adhered to throughout the procedure. All procedures were approved by the Animal Ethics Committee at Greater Bay Area Institute of Precision Medicine. Fecal samples were collected from two adult volunteers. The study was approved by the Institutional Review Board of Fudan University and written informed consent was obtained from either volunteer. The bacterial strains used for model training were previously isolated from human feces or saliva.

### Laser Induced Forward Transfer (LIFT) of bacterial cells

The equipment, PRECI SCS-R300 (Hooke Instruments, Ltd., China), was turned on and adjusted according to standards following the manufacturer’s handbook^25,26^. Samples with microbial cells were filtered and washed with sterile water, centrifuged at 3,000 rpm for 5 min, resuspended in sterile water, and spotted onto a slide (HSC24, Hooke Instruments, Ltd.). Visually selected bacterial cells (the manufacturer’s software) were LIFT-transferred into collectors (HSR04, Hooke Instruments, Ltd.) using a laser power of 80-120 nJ and then flipped into individual PCR tubes with brief spinning at 2000g (Yooning Mini-6K). The laser spot size was fixed, while the ejection size was adjustable (0.5-20 μm) via energy modulation to accommodate bacterial dimensions.

### Single-cell morphology

The morphologic features of microbial single cells include length, width, aspect ratio, area, perimeter, circularity, eccentricity, convexity, extent, and median centroid-contour distance. Convexity is the area of the contour divided by the area of the convex hull, while extent is the area of the masked region divided by the area of the minimum bounding rectangle. Microscopic images were analyzed using the Segment Anything Model^27^, which generated segmentation masks for each cell. These masks were then analyzed with OpenCV to measure the morphological features.

### Raman spectrometry

Raman spectra were collected using the PRECI SCS-R300, which is equipped with a 532 nm laser and a -70 °C cooled charge-coupled device detector with a 600 l/mm optical grating. The Raman signal was collected in the spectral interval from 400 cm^-1^ to 3600 cm^-1^. For cultural purpose, the laser power was 1 mW with an acquisition time of 1 s. For reference products of biological molecules, the laser power was 10 mW with an acquisition time of 10 s. Pre-processing of the raw spectra included cosmic ray removal, noise reduction with the Savitzky-Golay filter, baseline correction with the Adaptive Iteratively Reweighted Penalized Least Squares (airPLS) algorithm, and min-max normalization. Spectra with signal-to-noise ratio less than two were excluded from the analysis. The normalized intensity of the spectra was converted to the integrated area of each Raman shift within ± 10 cm^-1^ using Simpson’s rule, which approximates the integral by fitting parabolas to segments of the sampled data points. For molecular quantification, spectral signal in the fingerprint region (600-1800 cm^-1^) was used. Non-negative least squares (NNLS) fitting was used to fit integrated areas from cell spectra with the integrated areas from multiple reference spectra. The resulting weights from the NNLS fitting were used as estimates for the quantification of the molecules. For taxonomic identification, spectral region from 600 cm^-1^ to 3000 cm^-1^ was used.

### Machine learning

We created a database of single-cell morphology and Raman spectra, complete with taxonomic labels, using strains that were previously isolated from healthy volunteers. Bacterial isolates were cultured in Gifu Anaerobic Medium Broth, Modified (HOPEBIO, HB8518) before measurement. The database included 25 bacterial species, with each species comprising an average of around 500 cells. The training set for the Siamese network consisted of pairs of single cells organized from 100 cells per taxon, either from the same taxon (positive pairs) or from different taxa (negative pairs). The deep neural network received as input a vector that concatenated 10 morphological features with 2401 Raman spectral features (600-3000 cm^-1^). The network architectures for extracting features from morphological and spectral data were adapted from the Feature Tokenizer (FT) Transformer^28^ and ResNet^6^, respectively, by replacing the prediction layers with layers that output feature vectors of dimension 128. Features vectors from morphology and spectra were concatenated to form a final feature vector of dimension 256. The models were trained using contrastive loss based on the Euclidean distance between the feature vectors of two cells. We used the Adam optimizer to train the weights, with a learning rate of 0.001. A batch size of 512 was used. The feature vectors of the cells used in training were extracted using the trained models and constituted a reference library for cell matching. Cells that were not used to train the Siamese network were utilized to train (80% of the cells) and test (20% of the cells) an elastic net model for a specific taxon, predicting whether each cell belonged to that taxon. The predictors of the elastic net model were the Euclidean distances between a cell and the 100 cells used to train the Siamese network. For cells in microbiome samples, feature vectors were extracted from their morphological and spectral data, and their distances to reference cells were input into the elastic net model to determine whether they belonged to the desired taxa.

### Bacterial cell culture

Single cells were transferred to culture tubes and grown under anaerobic conditions (5% H_2_, 10% CO_2_, and 85% N_2_) in an anaerobic chamber (Coy Laboratory, Type B) at 37°C. Bacterial cells from mouse amniotic fluid samples were cultured in Wilkins-Chalgren Anaerobe Broth (Solarbio, LA4420). Bacterial cells from human fecal samples were cultured in Gifu Anaerobic Medium Broth, Modified (HOPEBIO, HB8518).

### Whole-genome sequencing

Taxonomy of cultured bacteria was determined by whole-genome sequencing. DNA was extracted using the VAMNE Magnetic Pathogen DNA Kit (Vazyme, DM202-01) according to the manufacturer’s instructions. Paired-end libraries (2 × 150 bp) were prepared using the VAHTS Universal Plus DNA Library Prep Kit (Vazyme, ND617) and sequenced on the DNBSEQ-T7 sequencer (MGI). Sequencing reads were quality filtered using fastp^29^ v0.22.0 and assembled by MEGAHIT^30^ v1.2.9. GTDB-Tk^31^ v1.0.2 was used for genomic taxonomy classification. Whole-genome ANI was estimated using FastANI^32^ v1.33. Mash distance was generated using Mash^33^ v2.0. SNPs were called using Snippy v4.6.0 (https://github.com/tseemann/snippy) and SNP-based phylogeny was constructed using FastTree^34^ v2.1.11. The structures of three variants of PBP2 were initially predicted using NeuralPLexer^35^ v0.1.0 and then optimized using DynamicBind^36^ v1.0. The binding affinities were estimated by DynamicBind and docking poses were scored using the RTMScore function^37^.

### Processing of fecal samples

Fresh fecal samples were self-collected using 50 ml centrifuge tubes (Thermo Fisher, 339652PK) filled with nitrogen. A sterile, disposable 3 ml dropping pipette (Biosharp, DG-300) was used to core out a 1 g sample, which was transferred to a sterile 15 ml centrifuge tube (Thermo Fisher, 339650). This sample was then immediately transferred to an anaerobic chamber (LABIOPHY, AL-B), homogenized in 5 ml of pre-reduced PBS through thorough vortexing, and filtered through a 40 μm filter (NEST, 258369) to remove dietary debris.

## Acknowledgements

The authors were grateful to Zhong Yang, Jiong Ma, Xu Zhang, Hang Li, Meiya Lin for equipment support. Ping Sun, Miaoyi Kuang, Ying Li for sequencing support.

## Author contributions

H.J. conceived and designed the study. Q.L., X.L., J.W., W.W., and L.L. carried out all experiments. Q.L., X.L., and R.G. performed data analysis. X.T., G.Z., R.G., and H.J. supervised the study. Q.L. and H.J. drafted the manuscript. All authors read and approved the final version of the manuscript.

