## Supplementary Figures for "Precision culturomics enabled by unlabeled single-cell morphology and Raman spectra"

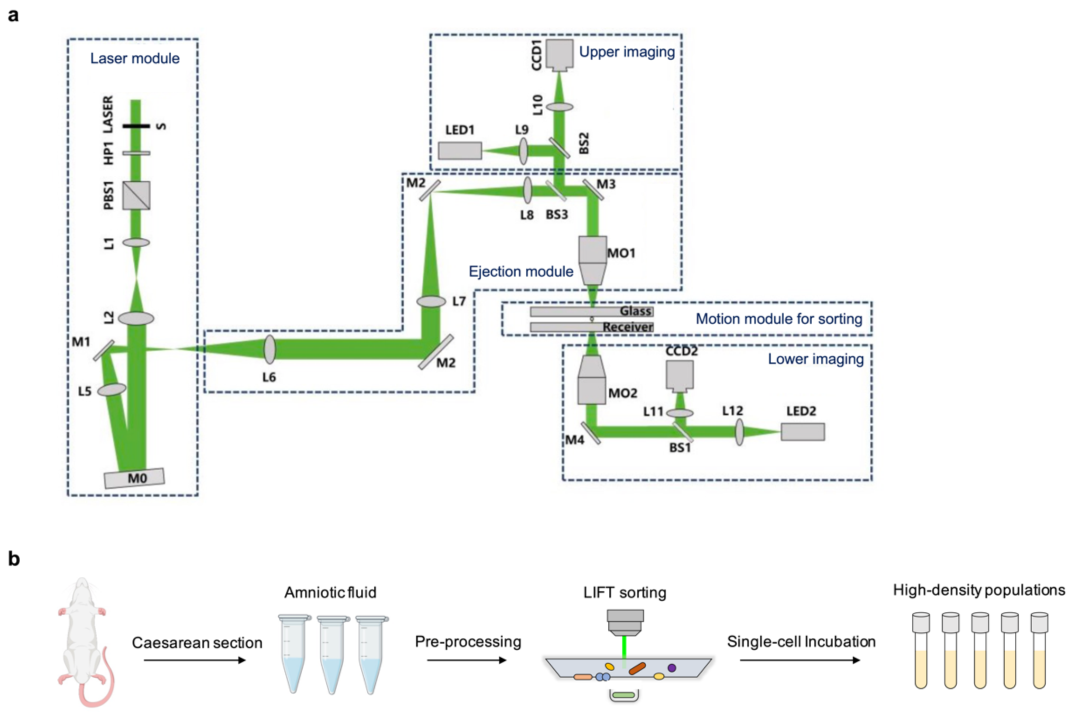


**Figure S1**. **Live culture from microliters of samples.**

(**a**) Optical setup for LIFT, morphology and Raman spectrometry (adapted from[26]). The commercially available PRECI SCS R300 device contains an adjustable, upright laser for LIFT, and an inverted laser for Raman spectra, both at 532 nm wavelength. (**b**) Schematic of microbial single-cell culturing from mouse amniotic fluid. Diagram created with BioGDP.com.

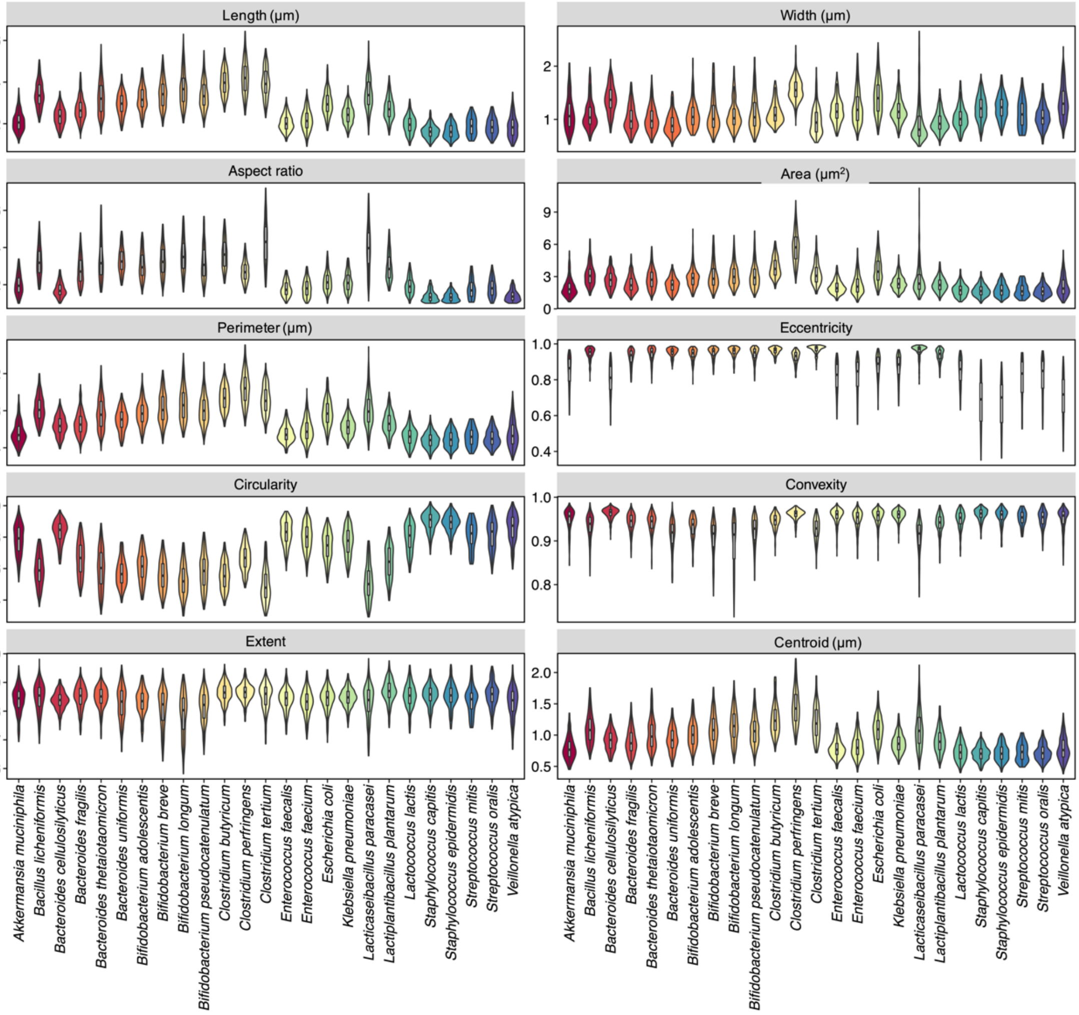


**Figure S2**. **Distribution of morphological features across microbial species.**
Violin plots illustrating the distribution of single-cell morphological features across different microbial species. Each violin represents the density of the data at various values, with wider sections indicating higher density and narrower sections indicating lower density. The boxplots within each violin denote the median and the interquartile range.

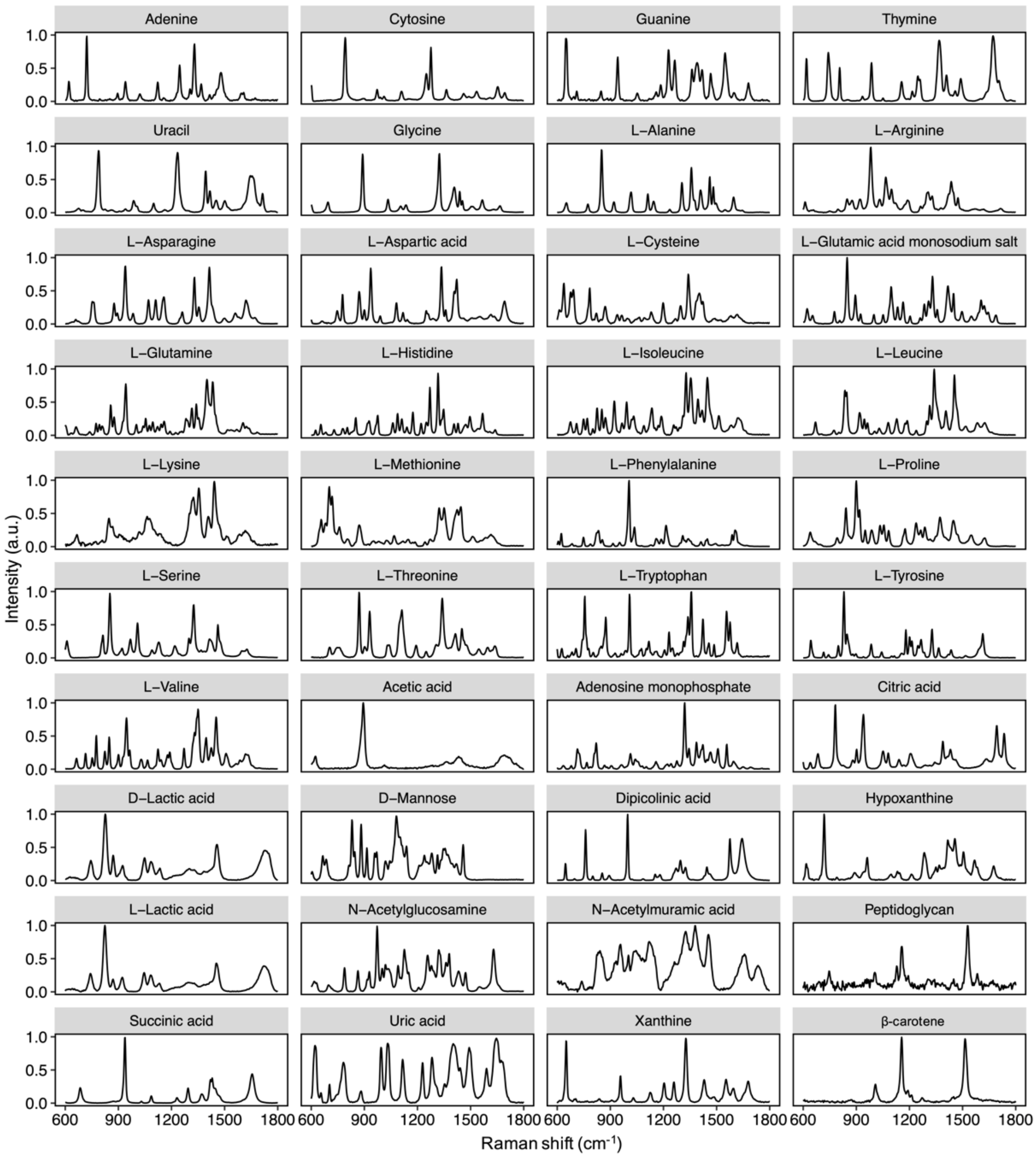


**Figure S3. Raman spectra of biological molecules used for quantification.**

Average Raman spectra of biological molecules used for molecule quantification for Raman spectra of bacterial cells.

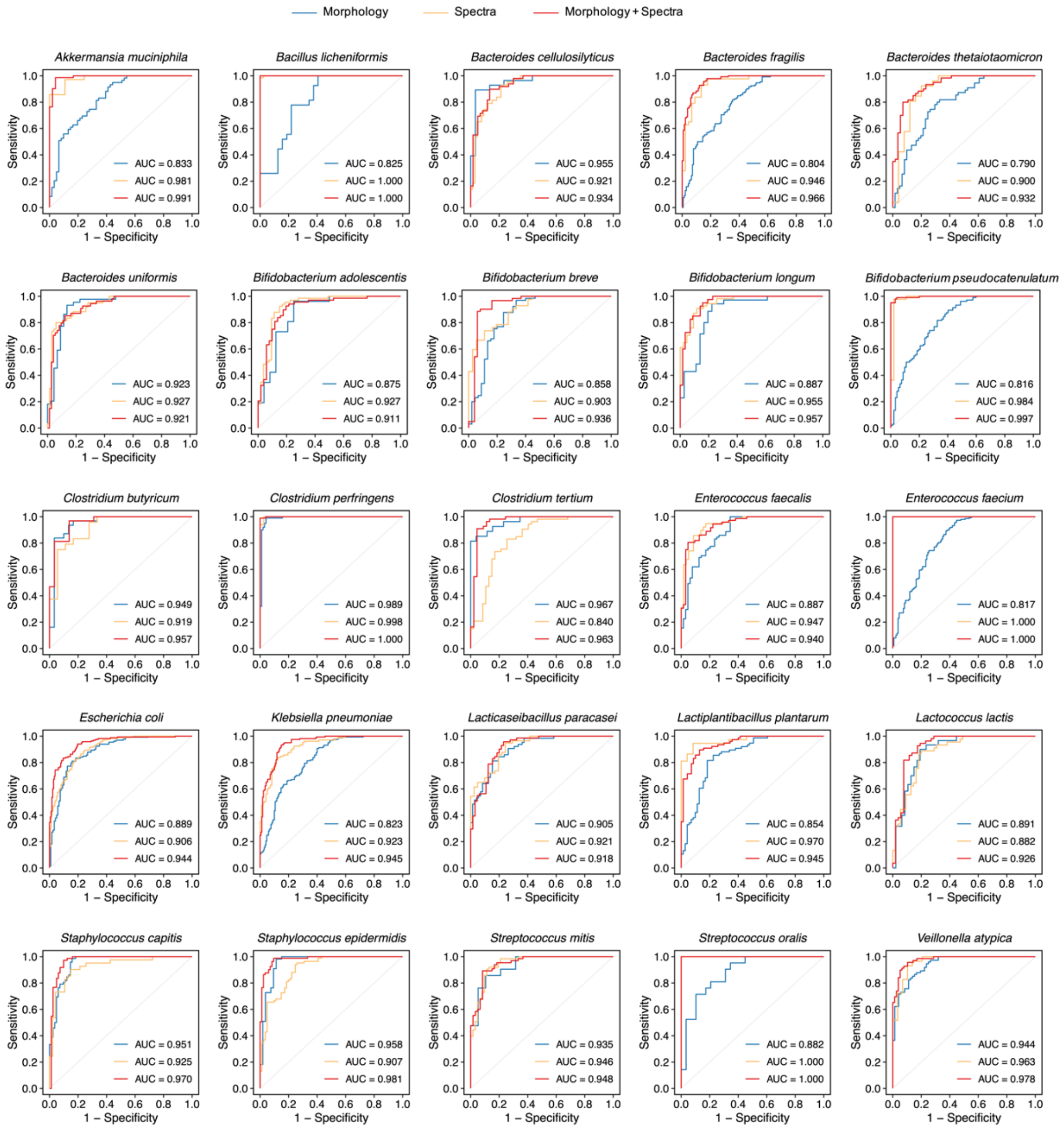


**Figure S4. ROC curves for elastic net predicting target species.**

Elastic net that predicts whether a cell belongs to a target species was created for each species based on morphology, Raman spectra, or a combination of both morphology and spectra.

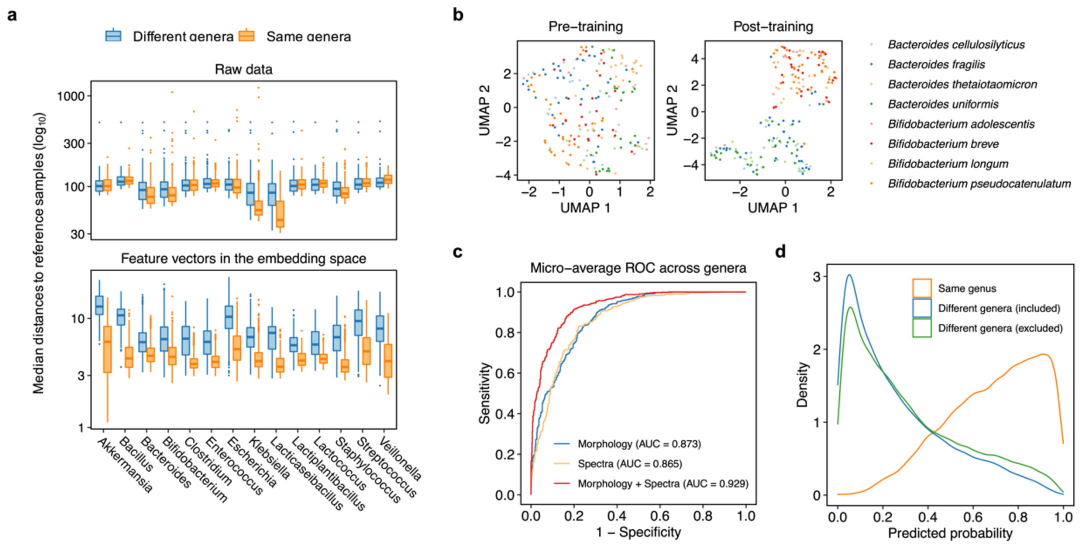


**Figure S5**. **The genus-level contrastive learning model.**

(**a**) Median Euclidean distances of raw data or feature vectors in an embedding space between test cells of different or the same genera and reference cells. (**b**) UMAP embeddings of features vectors before and after the training of the contrastive learning model. Cells from four *Bacteroides* species and three *Bifidobacterium* species were plotted to illustrate the clustering of cells based on genus labels. (**c**) ROC curves for elastic net predicting target genera based solely on morphology, Raman spectra, or a combination of both morphology and spectra. (**d**) Distribution of the predicted probabilities of being a target genus for test cells from the same or different genus that were included in or excluded from the training sets.

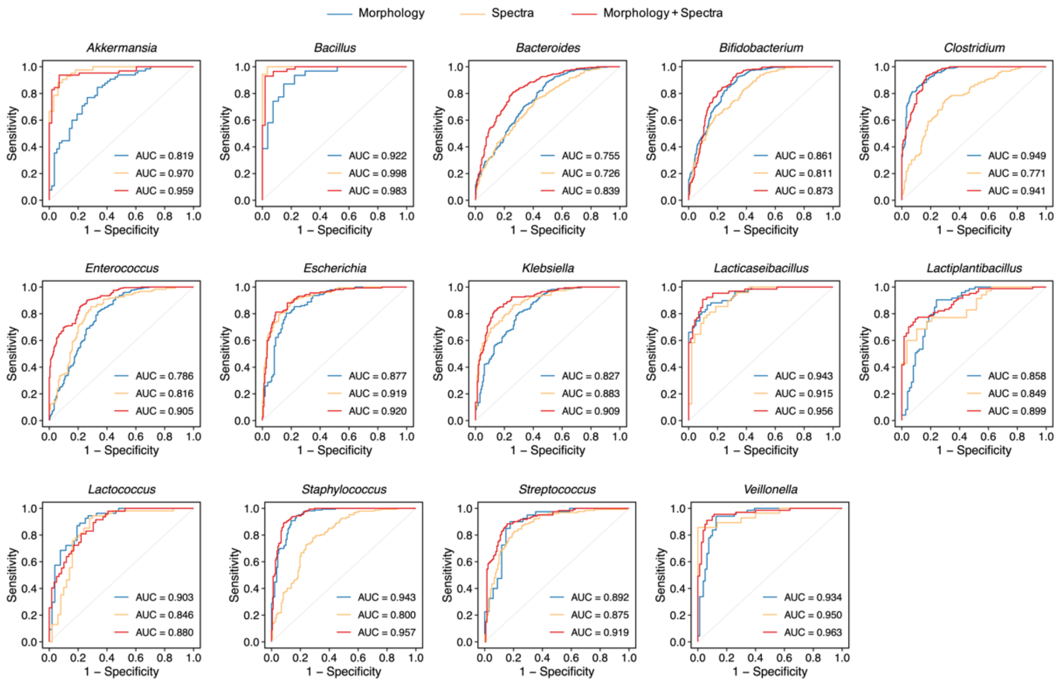


**Figure S6**. **ROC curves for elastic net predicting target genera.**

Elastic net that predicts whether a cell belongs to a target genus was created for each genus based on morphology, Raman spectra, or a combination of both morphology and spectra.

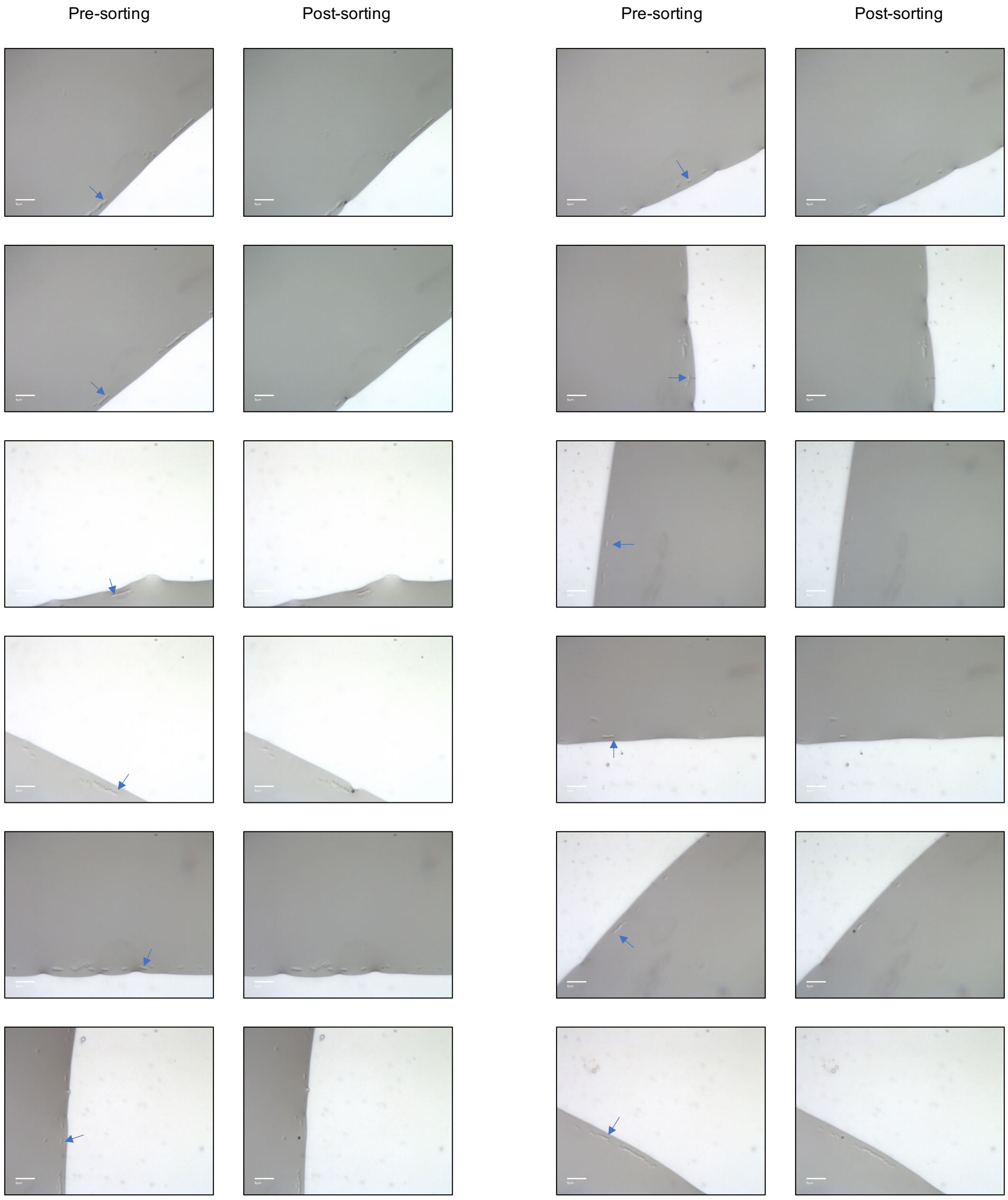


**Figure S7. Microscopic fields showing single-cell sorting from fecal samples.** Microscopic fields before and after sorting are shown, with blue arrows indicating the sorted cells.
